# Olfactory biopsy analysis of Alzheimer’s pathobiology across disease stages

**DOI:** 10.1101/2025.11.11.687356

**Authors:** Vincent M. D’Anniballe, Sarah Kim, John B. Finlay, Michael Wang, Tiffany Ko, Sheng Luo, Heather E. Whitson, Kim G. Johnson, Bradley J. Goldstein

## Abstract

Alzheimer’s Disease (AD) is a neurodegenerative condition affecting millions worldwide. Defining early pathobiological events remains challenging, in part due to inaccessibility of neural tissue. Because olfactory neurons are accessible, and olfactory loss is prevalent in AD, we evaluated olfactory brush biopsies from controls, individuals with cerebrospinal fluid (CSF) biomarker-confirmed AD, and cognitively typical individuals whose positive biomarkers signal a pre-clinical AD stage. We define via single cell RNA-sequencing (n=22 subjects) conserved neuroinflammatory T cell, myeloid, and olfactory neuron changes detectable even in pre-clinical AD subjects. Activated memory T cell states were a hallmark of pre-clinical AD, paralleling CSF T cell phenotypes seen in advanced disease, accompanied by both microglia-like inflammatory programs and olfactory neuron inflammatory injury. Together, our findings establish a novel platform permitting analysis of neural tissue in AD at its earliest stages.

## Main Text

Anosmia, loss of the sense of olfaction, is strongly associated with the development of Alzheimer’s Disease (AD), yet mechanisms driving olfactory loss in AD, and its relationship to disease progression, remain unclear. Positioned in the superior nasal cavity, the olfactory epithelium (OE) harbors olfactory sensory neurons (OSNs) whose axons project to glomeruli in the olfactory bulb, where they synapse with mitral/tufted cells that send second-order projections to primary olfactory cortices, including portions of the entorhinal cortex—one of the earliest brain regions compromised in AD^1^. Similar to the AD brain parenchyma, OSNs have been shown to accumulate amyloid-β plaques and neurofibrillary tau tangles, suggesting that the OE may faithfully mirror central AD neuropathology^2, 3, 4^. Moreover, because lymphatic drainage from the cerebrospinal fluid (CSF) traverses the cribriform plate ^5^, the OE may offer a readily accessible site to monitor the neuroinflammatory processes that are characteristic of AD.

Histologic studies of biopsies from the olfactory cleft, the olfactory sensory region of the nasal cavity, have uncovered local cellular changes linked to neurodegeneration-associated anosmia^4, 6, 7, 8^. Most of these assays relied on cadaveric tissue or samples from patients with advanced disease, leaving earlier stages underexplored^9, 10, 11^. Recent advances in single-cell transcriptomic profiling and minimally invasive, endoscopically guided cytology brush biopsies now enable high-resolution analysis of the local cellular milieu in specimens collected during routine outpatient visits at any disease stage. Therefore, we hypothesized that olfactory biopsies would offer a practical surrogate for nervous system tissue, facilitating identification of early pathogenic events in AD.

Here, we profiled OE clinic brush biopsies from three cohorts defined according to the revised 2024 Alzheimer’s Association’s biomarker-based diagnostic criteria^12^: cognitively typical adults with typical Aβ42/Aβ40 CSF levels (Controls), adults with atypical cognition and atypical Aβ42/Aβ40 levels (Clinical AD), and cognitively typical adults with atypical Aβ42/Aβ40 levels (Pre-Clinical AD) who are likely to progress to Clinical AD. By integrating neuronal and immune signatures across cohorts, we identified molecular shifts that track disease stage and yield tractable targets for therapeutic or diagnostic exploration. Ultimately, these findings position OE sampling as a novel practical strategy for studying AD pathobiology at its earliest, most treatable stage.

### Clinic brush biopsies for sampling cells of the olfactory cleft

The cell bodies of olfactory neurons are situated in the surface epithelium lining the airway of the nasal cavity superiorly, a region called the olfactory cleft (Fig. 1A,B). These olfactory sensory neurons (OSNs) detect inspired odors and project intracranially to synapse in the olfactory bulbs of the brain. As such, the neurons and associated cells within the OE are accessible via visualization using a nasal endoscope and can be sampled for biopsy using a small cytology brush in awake subjects (Fig. 1C, Supplementary Video 1). Live cell suspensions may be prepared for single cell RNA-sequencing (scRNA-seq), flow cytometry, or other assays^12–14^. Visualization of scRNA-seq from subjects with AD or controls biopsied for this study (n=22 subjects) demonstrates capture of OE cells and associated local immune populations, including T cells, macrophages, dendritic cells (DC), sustentacular cells, microvillar cells (MV), and OSNs (Fig. 1D-F, Extended Data Fig 1). The brush biopsy approach reliably yields olfactory cells, as well as variable amounts of adjacent surface respiratory epithelium, permitting downstream analysis of a combined dataset comprised of 220,509 cells, without contamination by deeper stromal, vascular, or systemic circulating immune cells.

**Figure 1.**
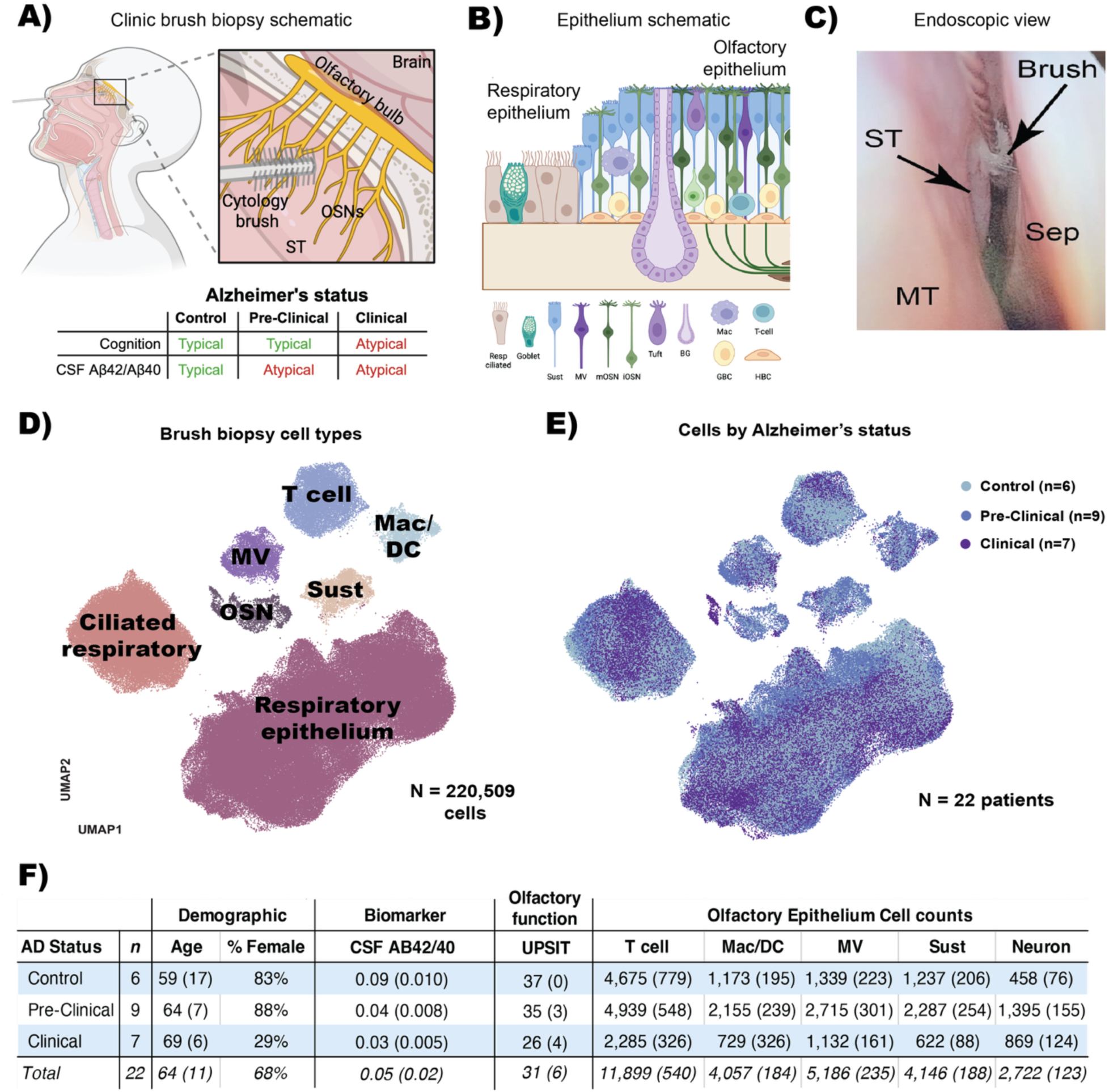
Brush biopsies as a practical strategy for sampling olfactory epithelium in Alzheimer’s Disease. **A)** Schematic showing brush biopsy of the olfactory cleft in Control (typical cognition and CSF biomarkers), Pre-clinical AD (typical cognition and atypical CSF biomarkers) and Clinical AD (atypical cognition and CSF biomarkers) subjects. ST = superior turbinate. **B)** Schematic showing the principal cell types that are sampled via olfactory cleft brush biopsy. Sust = sustentacular; MV = microvillar; mOSN = mature olfactory sensory neuron; iOSN = immature olfactory sensory neuron; BG = bowman’s gland; Mac = macrophage; DC = dendritic cell; GBC = globose basal cell; HBC = horizontal basal cell. **C)** zero-degree endoscopic view of brush biopsy technique. ST = superior turbinate; MT = middle turbinate; Sep = nasal septum. **D)** UMAP shows brush biopsy cell types identified by single cell RNA-sequencing. **E)** same UMAP by Alzheimer’s status. **F)** Table showing demographic, biomarker and risk factor variables, and olfactory epithelium cell counts by Alzheimer’s status. Demographic and biomarker variables are shown with average (SD) per row. Cell counts are total per cell type (columns) per Alzheimer’s status (rows) with average per subject per status in parentheses. (n = 22 subjects). UPSIT = University of Pennsylvania Smell Identification Test.

### The olfactory epithelium parallels CSF T-cell inflammation in pre-clinical AD

The CSF serves as a conduit for central nervous system antigens to be sampled by the peripheral immune system. This exchange occurs in part by CSF drainage through the lymphatic nasopharyngeal plexus—which transverses the OE—before reaching the deep cervical lymph nodes ^5, 13^. Recently, encephalitogenic CD8 memory T cells have been identified patrolling the CSF in individuals with clinical AD ^14^. Given the OE’s direct anatomical interface with this CSF drainage pathway, we hypothesized that a similar inflammatory T cell profile may also be present in the OE of AD subjects. To investigate this, we integrated single-cell transcriptomic data from 22 newly obtained OE biopsies (Control: n = 6, Pre-Clinical AD: n = 9, Clinical AD: n = 7) with published CSF datasets (Control: n = 9, Clinical AD: n = 9) to compare T cell subtypes and their function across the AD continuum.

Conventional scRNA-seq clustering approaches, which assign cells to discrete categories, can obscure the continuous and overlapping nature of T cell transcriptional programs. To overcome this, we used previously published and experimentally validated gene expression programs (GEPs), derived from consensus non-negative matrix factorization (cNMF) of 1.7 million T cells from 700 individuals across 38 tissues, to model each T cell as a mixture of functional and identity-defining signals^15^. Projecting the validated GEPs onto OE and CSF T cells was preceded by cross-cohort harmonization and batch correction (see Methods for details), enabling reproducible characterization and unsupervised cross-tissue comparisons of 20,039 CSF T cells and 10,678 OE T cells (Fig. 2A-C, Extended Data Fig 2A).

**Figure 2.**
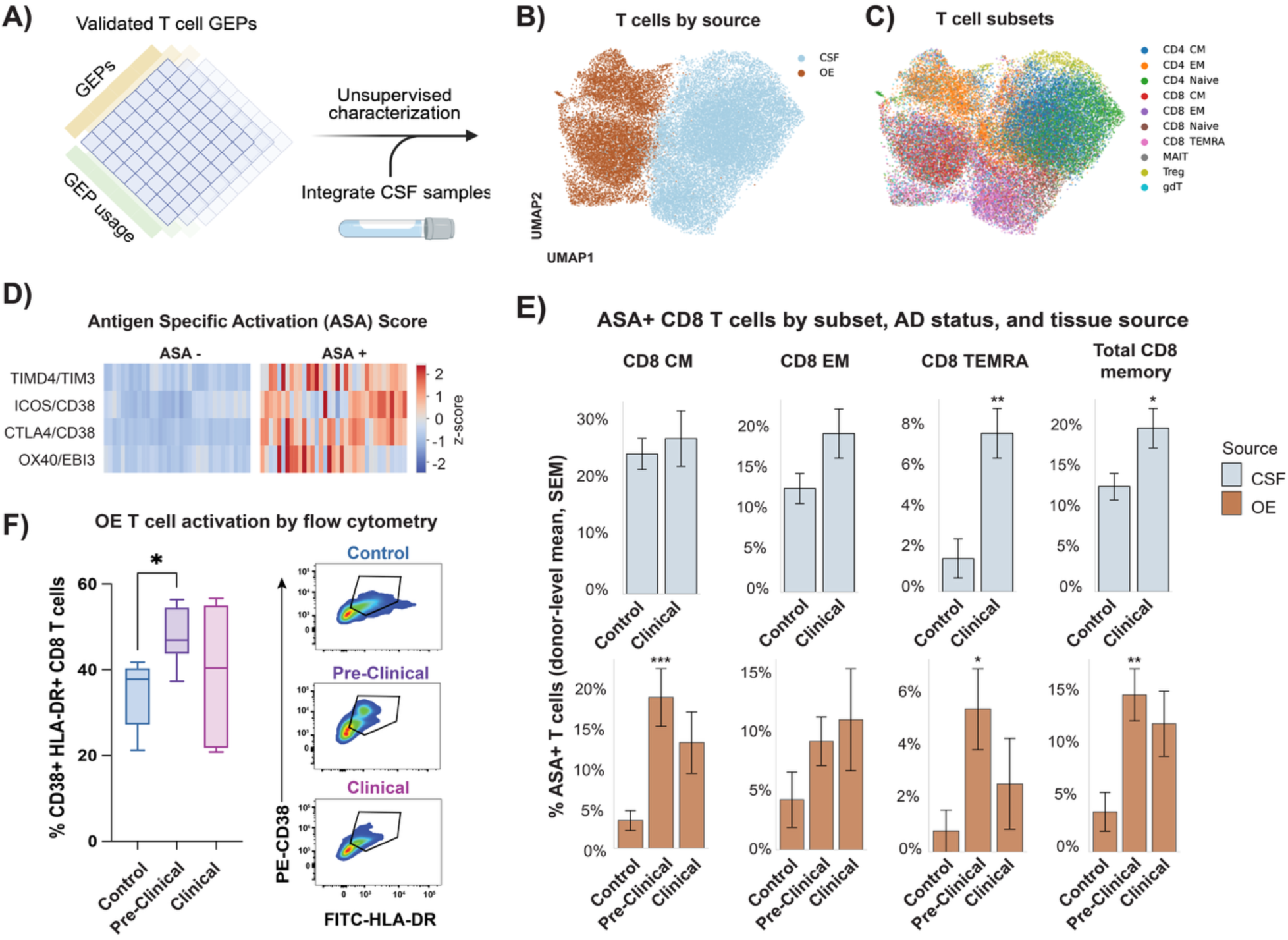
Olfactory epithelium parallels clinical AD CSF T-cell inflammation at the pre-clinical stage. **A)** Olfactory epithelium brush biopsies were obtained from Control (n=5), Pre-Clinical AD (n=9), and Clinical AD (n=5) subjects in clinic, and processed for scRNA-seq. Experimentally validated gene expression programs (GEPs) were projected onto T cells from each condition, and published CSF samples from AD subjects were integrated for comparison. **B)** UMAP of T cells by source (CSF or OE) and **C)** T cell subsets (CM = Central memory; EM = Effector memory; TEMRA = Terminal effector memory CD45RA; mucosal-associated invariant T (MAIT) cells). **D)** Heatmaps display the antigen-specific activation (ASA) score used to quantify inflammatory changes in T cells; each row represents a GEP previously shown to predict TCR signaling, while each column corresponds to a pseudobulked subject sample from the OE or CSF, comparing Z-scored expression between ASA-positive and ASA-negative cells. **E)** Bar graphs show the mean ± SEM of the percent ASA⁺ CD8 T cells across donor-level pseudo-bulk replicates (30-cell replicates per donor), stratified by CD8 subset, AD status, and source. Statistical comparisons were performed using a binomial generalized linear model (logit link) with Huber-White standard errors clustered by donor. * p < 0.05, ** p < 0.01, *** p < 0.001. **F)** Box and whisker plots show flow cytometry data of percent double-positive CD38 and HLA-DR CD8 T cells from the OE of Control (n = 3 subjects), Pre-Clinical (n=3 subjects), and Clinical (n=3 subjects) AD stages. Boxes indicate the median and interquartile range (25th–75th percentiles), and whiskers extend to 1.5x the interquartile range. Pseudocolor plots show representative CD38, HLA-DR double positive CD8 T cells from each condition. * p < 0.05, one way ANOVA with Tukey’s multiple comparisons test.

Except for mucosal-associated invariant T (MAIT) cells, the OE and CSF shared similar T cell subsets, while the OE contained a greater proportion of total memory CD8 T cells (OE = 41%, CSF = 21% p < 0.001), and a lower CD4/CD8 ratio (OE = 0.40, CSF = 0.70, p < 0.001) compared to CSF (Extended Data Fig 2B,C). Using an experimentally validated antigen-specific activation (ASA) score comprised of 4 T-cell function GEPs (labeled TIMD4/TIM3, ICOS/CD38, CTLA4/CD38, OX40/EBI3; Fig 2D), our modeling found that total CSF memory CD8 T cells contained significantly higher ASA in Clinical AD vs Control groups (20.1% vs 13.3%, p < 0.01, Fig 2E), consistent across several CD8 memory T-cell subsets. Strikingly, T cells from the OE paralleled CSF activation across the same T cell subsets, but was detectable at the earlier Pre-clinical AD stage compared to Controls (13.9 vs 3.5%, p < 0.001 Fig 2E). To validate these findings at the protein level, we quantified CD38+ HLA-DR+ CD8+ T cells by flow cytometry, an established phenotype of antigen-driven T cell receptor (TCR) activation and again found heightened inflammatory signatures in OE CD8 T cells during the pre-clinical AD stage compared to controls (p < 0.05, Fig 2F, Extended Data Fig 2D,E). These findings suggest that T cell activation within the OE could emerge early in AD, and parallel immune alterations detected in the CSF.

### Myeloid lineage cells contribute to AD-related OE inflammation

The OE is a mucosal barrier with a dense resident myeloid compartment that supports neuro-immune surveillance^17^. In our dataset, the majority of OE myeloid clusters expressed the purinergic receptor P2RY13 (Extended Data Fig. 2A-D), a microglial receptor linked to neurite-contact/surveillance^18^, supporting the view that neuron-interacting potential is distributed across multiple myeloid subsets in the olfactory neuroepithelium. Consistent with this biology, both human and animal studies have identified specialized macrophages in the olfactory mucosa, including populations with microglia-like features^19^. Other work shows that interactions between myeloid cells and T cells can promote tau-related neurodegeneration^20^, suggesting that OE myeloid populations could participate in broader AD-associated inflammatory pathways^21^. Accordingly, we hypothesized that neuron-interacting myeloid programs would be detectable in the human OE.

Guided by this rationale, we profiled the entire OE myeloid compartment by single-cell RNA-seq and applied a myeloid subset-agnostic cNMF to infer shared myeloid functional programs and their per-cell usage (Fig 3A,B, Extended Data Fig D-H). To avoid confounding by any single individual, program positivity was quantified at the donor level. GEPs for identity and function were drawn from a previously published, experimentally validated reference of 91,523 human myeloid cells encompassing microglial and inflammatory states learned in an unsupervised framework^22^. These reference GEPs were projected onto 3,482 OE myeloid cells, and cell identities (microglia-like, macrophages, and dendritic cells) were assigned based on dominant identity-program loadings. Baseline proportions of each myeloid identity were comparable across conditions (Extended Data Fig. 3C). Functional profiling then focused on four validated programs across subsets—complement-immunosuppressive, scavenger-immunosuppressive, systemic-inflammatory, and inflammatory-microglial—at single-cell resolution (Fig. 3B). This design captures functional states that can span manually labeled cell identities, while enabling fair, donor-level comparisons across clinical stages.

**Figure 3.**
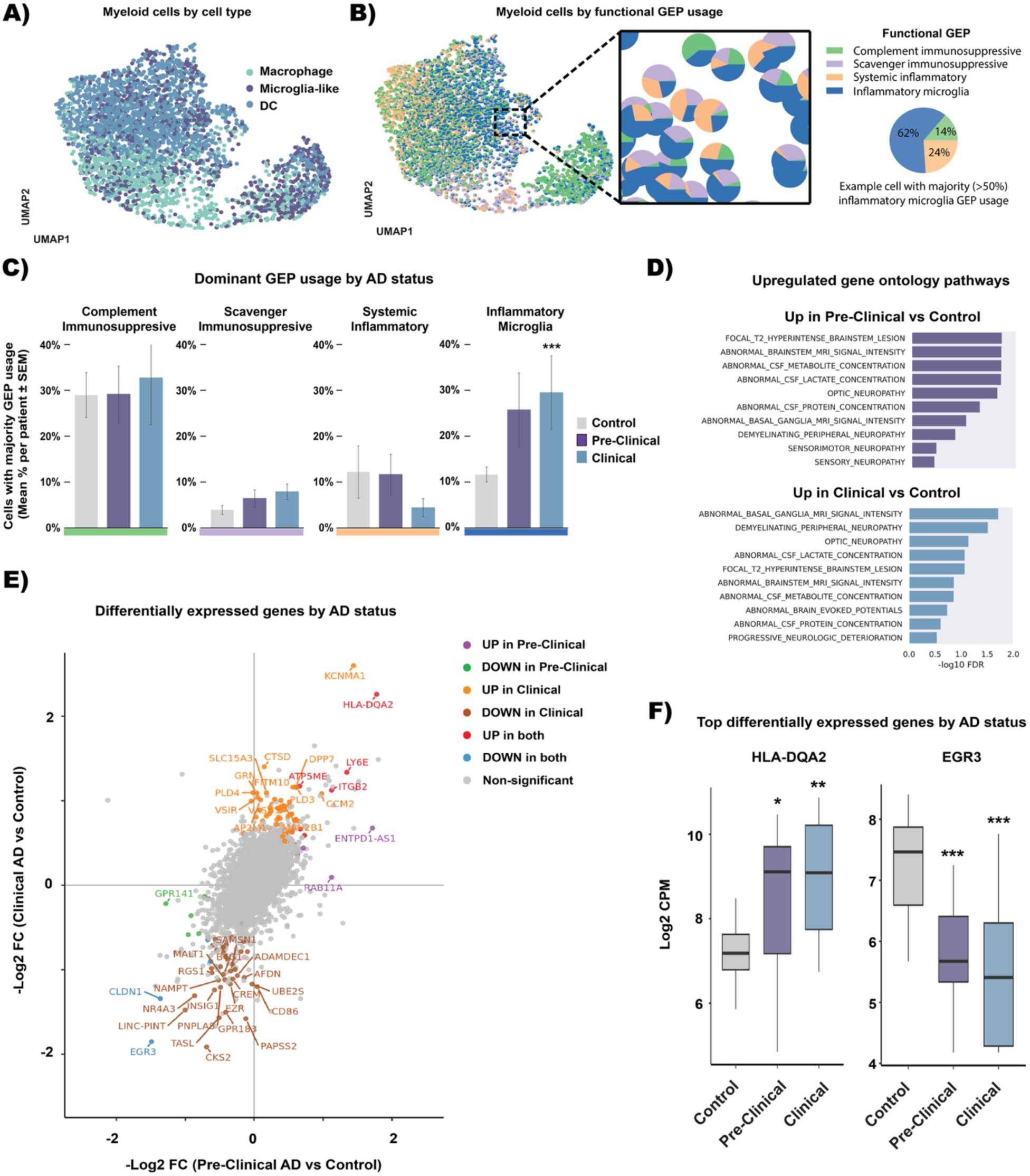
Myeloid cells in the OE demonstrate inflammatory changes associated with AD. **A)** UMAP shows the 4 principle myeloid cells in the OE, microglia-like macrophages, macrophages, monocytes, and conventional dendritic cells (cDC). **B**) Same UMAP as A) but with each cell as a pie chart showing the proportion of 4 functional myeloid GEPs, complement immunosuppressive, scavenger immunosuppressive, systemic inflammatory, and inflammatory microglia. Example cell pie chart on the right shows a cell with majority (>50% usage) of the functional inflammatory microglia GEP that were analyzed in C). **C**) Bar charts show the percentage of myeloid cells per subject (mean, SEM) that contain a majority (>50%) of either of the four functional GEPs by AD status. Kruskal–Wallis omnibus test followed by Bonferroni-corrected two-sided Mann–Whitney U tests for the three pairwise stage contrasts, ** p < 0.01. **D**) Top neuro-related up-regulated pathways (-log10 FDR) from fry enrichment in myeloid cells for Pre-Clinical (top) and Clinical AD (bottom) vs Control. **E**) Two-way volcano plot shows differentially expressed genes between Pre-Clinical AD vs. Control and Clinical AD vs. Control from subject-level pseudobulk differential expression analysis. Genes that pass FDR < 0.05 in either contrast are colored by which quadrant they fall into; genes that reach -Log FC ≥ 1.2 in either contrast are annotated. **F**) Box-and-whisker plots show the median and inter-quartile range of log2 CPM values for each subject-level pseudobulk for the most upregulated (HLA-DQA2) and downregulated (EGR3) genes in both contrasts. * p < 0.05 ** p < 0.01, *** p < 0.001; Wilcoxon rank-sum tests (Benjamini–Hochberg adjusted). (n=22 subjects).

Across all myeloid cells, we observed a clear enrichment of inflammatory microglia functional programs in clinical AD. Notably, the same pattern was already evident in pre-clinical AD, with effect sizes that increased from pre-clinical to clinical disease (Fig 3C). Moreover, subject-level analysis revealed marked similarities in human phenotype gene ontology terms between Pre-Clinical and Clinical AD compared to Controls, many of which are related to abnormal CSF metabolites and lesions of the brain (Fig 3D). Pseudobulk differential expression analysis revealed a concordant set of 15 genes whose expression was significantly increased or decreased in both the pre-clinical and clinical stages relative to controls (Fig 3E, Supplementary Data Tables S1, S2). For example, HLA-DQA2, the most strongly up-regulated gene in both pre-clinical and clinical AD, lies within the highly polymorphic HLA class II region on chromosome 6—an area also flagged by AD GWAS analyses ^23^—and, together with the tightly linked HLA-DR and HLA-DQ genes, encodes peptide-binding molecules that determine antigen presentation and subsequent T cell response to self- and foreign antigens ^24^. EGR3, a transcription factor essential for short-term memory and regulation of the synaptic-vesicle cycle, showed the strongest decrease in expression in both pre-clinical and clinical Alzheimer’s stages relative to controls ^25, 26^ (Fig 3F). Collectively, our data indicate that myeloid-cell alterations in the OE may intensify with AD progression and, crucially, are detectable during the asymptomatic pre-clinical phase.

### Olfactory sensory neurons demonstrate molecular and immune crosstalk changes in AD

The OSNs situated in the OE, whose axons feed to the olfactory bulbs, are replaced by basal stem and progenitor cells throughout life to maintain function ^9, 27^. Thus, the peripheral olfactory system harbors neurogenesis, neuronal maturation, axon growth and targeting, and synaptogenesis continuously, potentially making these afferent pathways especially susceptible to numerous pathologic mechanisms. Autopsy studies demonstrated that OE mirrors central AD neuropathology ^2, 7^, but no studies have reported scRNA-seq analysis from live human OE biopsies from AD subjects. Analyzing neuron lineages from our samples (n=7 AD; 9 Pre-clinical AD; 6 Control), we classified cells based on canonical genes that represent immediate neural precursor (INP), immature-(iOSN), or mature olfactory sensory neurons (mOSN) (Fig 4A,B) ^9^. Patient-level differential expression revealed extensive transcriptional changes in OSNs from both pre-clinical and clinical AD versus controls, with 8 genes displaying parallel alterations across disease stages (Fig 4C,D, Supplementary Data Tables S3, S4). For example, AKR1B1, which was highest in both pre-clinical and clinical AD OSNs relative to controls, encodes aldose reductase—the NADPH-consuming first enzyme of the polyol pathway—and also exhibits prostaglandin F2α synthase activity; both mechanisms can amplify oxidative and inflammatory signaling, and aldose reductase activity has been shown to regulate Aβ-induced microglial activation, making AKR1B1 up-regulation a plausible contributor to neuroimmune crosstalk in the OE^28^. In contrast, CERT1, which was lowest in both AD groups, encodes the ceramide transfer protein (CERT/CERTL) that shuttles ceramide from ER to Golgi for sphingomyelin synthesis; experimental augmentation of CERTL reduces brain C16-ceramide, Aβ burden, and neuroinflammation in AD models, and human CERT1 variants disrupt neuronal sphingolipid homeostasis—together supporting that reduced CERT1 may bias OSNs toward pro-amyloid, pro-inflammatory ceramide signaling^29^. Other differentially expressed AD transcripts of interest include enrichment of FIBP, which amplifies downstream FGF-signaling cascades and heightens microglial activation, positions it as a pivotal mediator of inflammatory neuron-to-myeloid crosstalk among AD subjects^30, 31^. Whereas the sustained depletion of LBH—previously observed in hippocampal neurons from post-mortem AD brains and inversely correlated with amyloid-β accumulation^32^—is similarly measurable in OSNs (Extended Data Fig 4).

**Figure 4.**
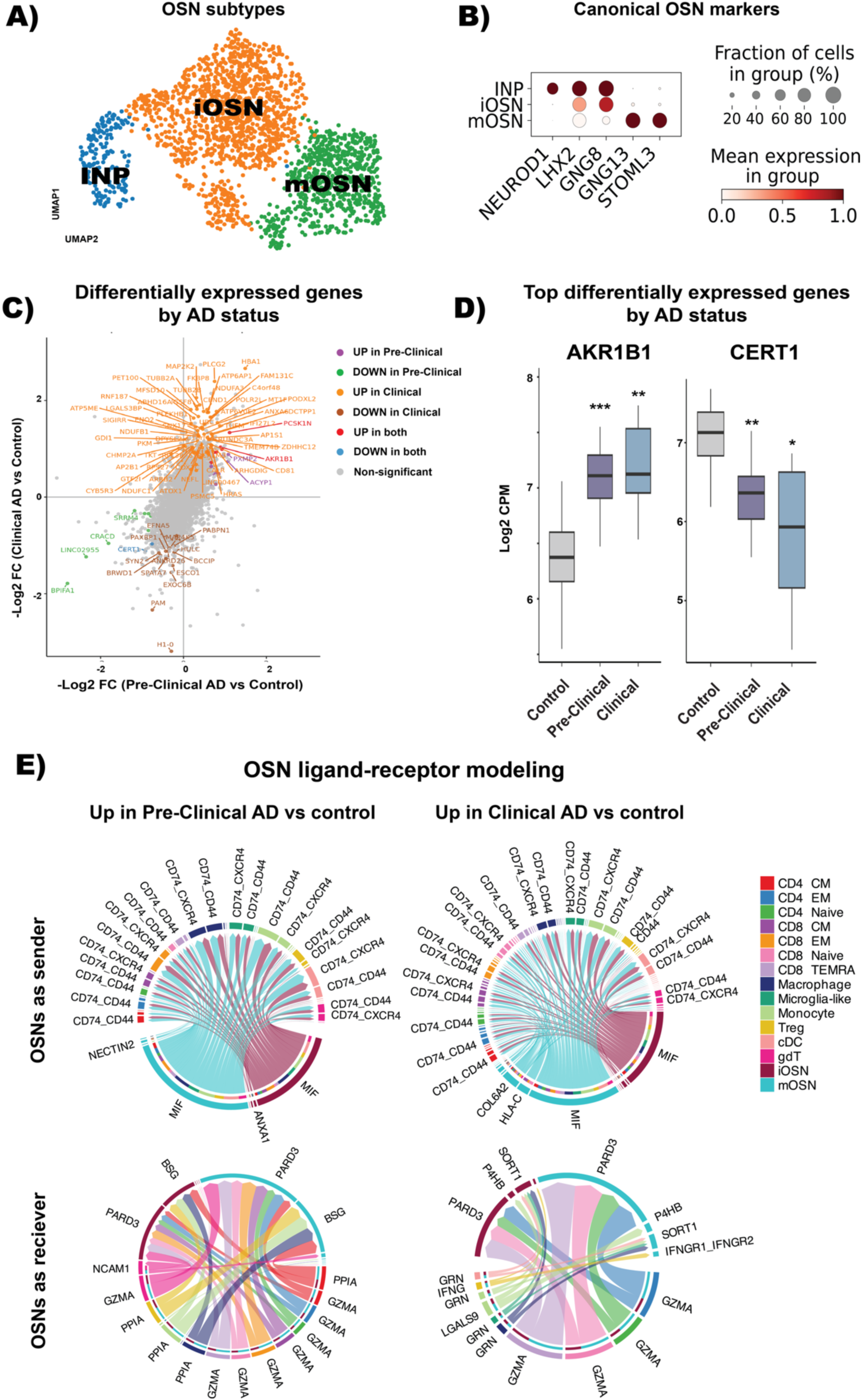
OSNs demonstrate AD pathology and heightened immune cell interaction potential. **A)** UMAP shows olfactory sensory neuron subsets. iOSN = immature olfactory sensory neuron; mOSN = mature olfactory sensory neuron; INP = immediate neuronal progenitor. **B)** Dot plot shows canonical gene profiles that were used to characterize each neuronal subset. **C)** Two-way volcano plot shows differentially expressed genes between Pre-Clinical AD vs. Control and Clinical AD vs. Control from subject-level pseudobulk differential expression analysis. Genes that pass FDR < 0.05 in either contrast are colored by which quadrant they fall into; genes that reach -Log FC ≥ 1.2 in either contrast are annotated. **D)** Box-and-whisker plots show the median and inter-quartile range of log2 CPM values for each subject-level pseudobulk for the most upregulated (AKR1B1) and downregulated (CERT1) genes in both contrasts. * p < 0.05 ** p < 0.01, *** p < 0.001; Wilcoxon rank-sum tests (Benjamini–Hochberg adjusted). **E)** Chord diagrams show ligand-receptor pathways that are differentially expressed in either Pre-Clinical (left) or Clinical AD (right) compared to controls with neurons as ligand sender to immune cells (top) or as ligand receivers from immune cells (bottom). (n = 22 subjects).

The robust olfactory T cell and myeloid inflammatory changes identified here at the Pre-Clinical AD stage, and evidence that experimental depletion of either immune population blocks tau-mediated neurodegeneration in murine models, suggests that OSNs may be especially vulnerable to inflammatory interactions between OSNs and immune cells, even pre-clinically. Modeling ligand-receptor interactions between OSNs and myeloid cells or T cells, potent differentially expressed interactions were predicted in neurons from AD subjects compared to Controls. For instance, pro-inflammatory signals such as granzyme A (GZMA), implicated in adaptive T cell mediated CNS pathobiology in AD, were identifiable in our OE samples at both disease stages (Fig 4E). Mirroring clinical AD, OSNs from pre-clinical AD subjects had markedly higher levels of macrophage migration inhibitory factor (MIF) signaling—a cytokine that colocalizes with Aβ plaques in APP mice, attenuates Aβ neurotoxicity when inhibited, and is elevated in the cerebrospinal fluid of human AD subjects ^33, 34^—toward nearly all immune subsets. Together, coordinated shifts in neuronal and immune cell states point to disrupted immune-neuron crosstalk in OE biopsies from both clinical and pre-clinical AD subjects.

### OE module scores provide a tissue-level readout of AD-related neuro-immune programs

Recent efforts to develop serum assays^35^ have led to FDA approval of the pTau217/ß-Amyloid 1-42 plasma ratio for detecting amyloid plaques in adults over 55, a remarkable progression in the field. While promising for pre-clinical AD diagnosis, the currently approved use of plasma assays remains limited to clinically manifest AD. These blood-based tests quantify biochemical end products of pathology—reflecting amyloid and tau accumulation—but have limited access to neural tissue and are therefore less able to resolve the upstream cellular processes that give rise to these changes. In contrast, the OE, a neural interface that itself accumulates hallmark AD pathology, may reveal transcriptional programs that evolve in parallel with or precede amyloid and tau deposition. We therefore asked whether an aggregate gene module derived from OE brushings could summarize AD-related programs at the tissue level, offering a readout of neuroimmune activity that may occur upstream or in parallel with amyloid and tau pathology.

To address this, we next examined whether AD-related changes were most detectable in the immune compartment, the neuronal compartment, or across the entire OE. First, we re-clustered each sample based on the major OE cell types most frequently sampled per subject. The immune compartment included T cells, macrophages, and DCs, while the neuronal/support compartment consisted of neurons, sustentacular cells, and microvillar cells (Fig 5A). Next, we performed subject-level pseudobulk DE analysis comparing each compartment in pre-clinical or clinical AD to control, and visualized gene expression by cell compartment across subjects, including additional clinical data such as olfactory function measured by University of Pennsylvania Smell Identification Test (UPSIT) scores and ApoE allele status (Fig 5B). We found that using the difference in Z-score (ΔZ-score), either the olfactory or immune compartment yielded significant differences between Control and Pre-Clinical AD or Clinical AD groups (Fig 5C, Supplementary Data Tables 5 and 6). Moreover, the combined (i.e., immune and neuronal) OE module distinguished AD-associated groups with an AUC of 0.81 (95% CI: 0.68-0.94) (Fig 5D). The module correlated significantly with the CSF Aβ42/Aβ40 ratio (R = −0.57, p = 0.01) but showed no association with CSF p-Tau-181 or MoCA scores, suggesting that these genes could reflect early, amyloid-linked molecular changes in the OE rather than overall cognitive status (Fig 5E). Taken together, these results indicate that OE brushings capture coordinated neuronal and immune transcriptional shifts that parallel central AD pathology, providing a framework to study early disease mechanisms in accessible human tissue.

**Figure 5.**
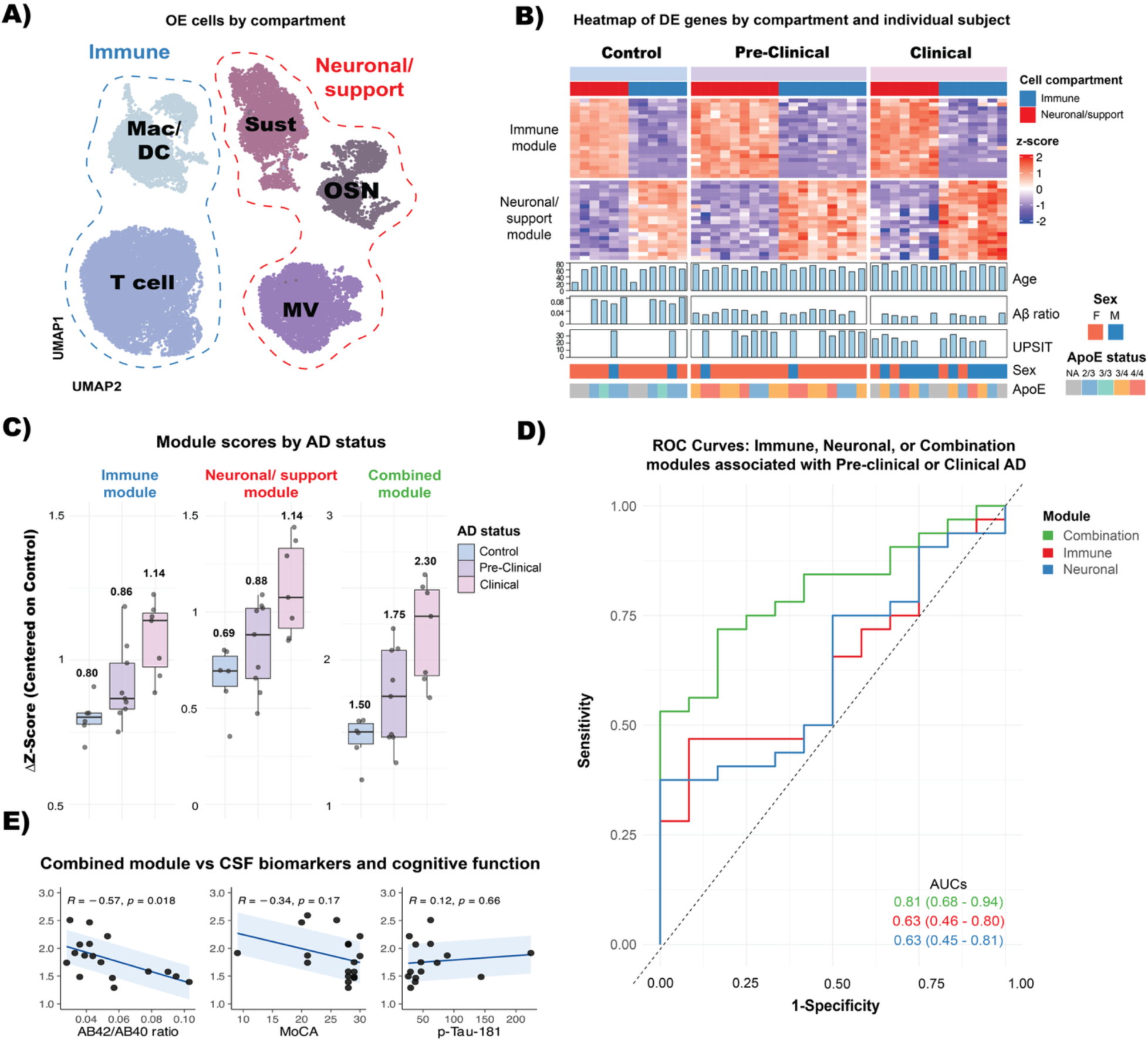
Module scores from whole OE brush biopsies associate with AD. **A)** UMAP shows five principal cell types sampled by olfactory cleft brush biopsies, categorized into neuronal support (MV, OSN, and Sustentacular) or immune (T cell, Macrophage, and DC) subsets. **B)** Heatmap shows the top differentially expressed genes by AD status and neuronal support or immune compartments. Each row represents a gene; each column represents an individual subject pseudobulk sample. Beneath the heatmap are demographic and biomarker variables for each individual subject including Age, Amyloid-beta ratio, UPSIT score, Sex, and APOE status. **C)** Box and whisker plots show the delta Z score of either the neuronal/support, immune, or combined module, centered on controls. **D)** shows receiver operating characteristic area under the curve (AUC) plots showing the sensitivity and specificity of either the immune module, neuronal support module, or combination in classifying Pre-clinical or Clinical AD compared to controls. AUC and its 95 % confidence interval were estimated from 2,000 stratified bootstrap replicates. **E)** Scatter plots show combined-module ΔZ versus the AB42/AB40 plasma ratio, MoCA score, and plasma p-Tau-181, with linear fits (blue), ±1 SD ribbons, and Pearson r and p values indicating the strength and significance of each association. (n = 22 subjects).

## Discussion

AD research is hampered by the impracticality of sampling live brain tissue, yet the OE, which harbors primary sensory neurons and an extensive immune milieu, may offer a window into central neurobiology. By generating a single-cell transcriptomic atlas from whole-brush biopsies of the olfactory cleft in 22 living subjects spanning controls, pre-clinical AD, and clinically manifest AD, we uncovered a continuum of disease-linked shifts detectable in biomarker-positive, cognitively unimpaired patients and mirrored in clinical AD. Activated CD8⁺ memory T cells and inflammatory myeloid programs were present in the pre-clinical group and were also present—often at greater magnitude—in the clinical group, while OSNs showed stress-related transcriptional signatures that overlap with those previously reported in postmortem AD brains. Together, these findings position minimally invasive OE sampling as a practical platform for studying early AD pathobiology.

Our analyses support the notion that an OE biopsy provides a stage-associated neural readout not readily obtained from other tissues. It samples a primary sensory neuroepithelium at a neuro-immune interface adjacent to CSF outflow, where OSNs and resident immune cells appear to register CNS-linked signals described in AD brains. As such, OE biopsy may provide tissue-level context on neuro-immune programs that bulk fluids such as plasma or cell-free CSF cannot resolve. Pending optimization and confirmation in larger studies, OE profiling could help clarify early AD pathobiology alongside amyloid and p-tau assays. One could envision the development of a targeted panel measurable from minimally processed brush biopsy material, assessed via longitudinal follow-up to assess prognostic value. The OE programs we report from 22 subjects align with AD brain literature and provide a concrete basis for focused follow-up.

Published olfactory mucosa histologic reports, from live or autopsy samples, identified classic hallmarks of AD pathology such as amyloid plaques and tauopathy, but these subjects had advanced disease, i.e. Braak stages V–VI^7^. More recent elegant AD single cell studies analyzing brain regions also necessarily relied upon cadaveric samples^36^. Here, we identify specific immune cell functional shifts in the OE from live subjects, accompanied by neuronal transcriptional alterations. These findings are consistent with a model in which not only the entorhinal cortex but also the peripheral olfactory organ are sites of AD pathobiology at early, even pre-clinical stages. Whether there is a causal relationship, or propagation of pathology along the afferent olfactory pathway, remains to be determined. For example, evidence has suggested that specific Herpesviruses may follow the OSN synapse at the olfactory bulb, to potentially propagate neuroinflammatory changes to the next node at the entorhinal cortex^37^. Regardless, early AD events clearly occur at the OE, providing a rationale to utilize OE brush biopsies from preclinical AD subjects to further define the earliest possible therapeutic targets aimed at arresting neuro-immune or other pathologic signals. Additionally, cross-compartment comparisons between OE and CSF were performed with matched T-cell populations and modern integration methods, yet cross-study differences in cohort composition, sequencing protocols, and diagnostic labels may still influence apparent concordance. We therefore interpret OE-CSF similarities as convergent patterns rather than proof of equivalence, and we do not infer causality across tissues. Definitive validation will require prospectively paired OE and CSF sampling in the same individuals with orthogonal protein and functional assays.

Olfactory cleft biopsies, even performed by highly experienced surgeons, have highly variable rates of success in capturing large numbers of neurons^9, 10^. This is likely due to several factors, including local anatomic variations, known variable patters of surface olfactory epithelial replacement by respiratory-like metaplasia^8, 11^, and challenges with selective viability issues with neurons in cell suspension. Accordingly, per-patient yields and cell-type fractions were transparently reported to make variability explicit (Fig. 1E). Most importantly, we intentionally designed our analyses to focus on within–cell-type transcriptional contrasts rather than proportional cell-type abundance. This approach allows biological inference even when total or cell-type-specific yields vary by biopsy. In doing so, we avoid composition-based claims that could conflate sampling variance with disease-linked differences, instead emphasizing conserved, cell-intrinsic transcriptional programs as the most reliable indicators of AD-related biology. Moreover, we base our analyses on publicly available human gene-expression programs validated across hundreds of subjects, ensuring a fully reproducible workflow that will effectively scale as more olfactory-biopsy datasets are generated. The single-cell atlas described here could also serve as a reference for future deconvolution of bulk RNA-seq datasets to infer cell-type-resolved signals independent of biopsy composition. As an immune barrier defense structure, we have found that olfactory epithelial immune cell populations are reliably captured in olfactory brush biopsies. Future studies may wish to use 5’ V(D)J-enabled reagents to couple transcriptomes with TCR sequences, enabling T cell clonotype analyses across AD stages.

Endoscopic olfactory-cleft brush biopsy could provide a feasible and repeatable way to sample accessible neuroepithelium in living humans at any disease stage. In our cohort, single-cell profiling of immune and OSN programs in OE revealed transcriptional states that track with biomarker-defined stages of AD. Larger, longitudinal studies that integrate OE profiling with CSF and plasma biomarkers, amyloid and tau PET, and clinical outcomes are needed to determine prognostic value and mechanism. If these findings are confirmed, OE profiling could complement established modalities by clarifying early pathobiology and by informing the selection and monitoring of candidates for disease-modifying therapies.

## Author contributions

Conceptualization: BJG, HEW

Methodology: VD, SK, JF, MW, TK, SL, KGJ, BJG

Biopsies: BJG

Visualization: VD, BJG

Funding acquisition: BJG, HEW

Project administration: BJG, KGJ

Supervision: HEW, KGJ, SL, BJG

Writing – original draft: VD, BJG

Writing – review & editing: VD, HEW, SL, KGJ, BJG

## Data and materials availability

scRNA-seq raw and processed data are available at GEO record GSE302937

## Methods

### Human participants

This research was performed under protocols approved by the Institutional Review Board of Duke University under IRB protocols 00088414 and 00116591. Biopsy subjects were recruited from participants enrolled in the Duke University and the University of North Carolina Alzheimer’s Disease Research Center (ADRC). The ADRC collected biomarker data including cerebrospinal fluid (CSF) Aβ42/Aβ40 ratio, APOE genotyping, and University of Pennsylvania Smell Identification Test (UPSIT). The UPSIT is a validated 40-item test of olfactory function ^1^. Alzheimer’s Disease (AD) status for each subject was classified according to the revised 2024 Alzheimer’s Association’s biomarker-based diagnostic criteria: cognitively typical adults with typical Aβ42/Aβ40 CSF levels (Controls), cognitively typical adults with atypical Aβ42/Aβ40 levels (Pre-Clinical AD), and adults with atypical cognition and atypical Aβ42/Aβ40 levels (Clinical AD). All laboratory members, including the performing surgeon, were blinded to AD status at time of tissue collection. No randomization was performed, all subjects were selected based on their AD status. Participants were selected at random from ADRC enrollees who had consented to research biopsy. All eligible individuals were approached, minimizing self-selection bias; recruitment pathway is not expected to influence molecular outcomes.

### Olfactory mucosa brush biopsies

All olfactory epithelial biopsies were performed by a board-certified otolaryngologist (BJG) in the Duke Department of Head and Neck Surgery & Communication Sciences Rhinology clinics using cytology brush biopsy in awake subjects, following informed consent. Topical nasal oxymetazoline/tetracaine spray was used as a local anesthetic. Rigid nasal endoscopy was performed to exclude evidence of sinonasal disease such as mass, edema, nasal polyps, or infection. One side was chosen for biopsy, based on best visualization of the olfactory cleft region superiorly and posteriorly, and biopsy was performed by gently positioning a cytology brush (Cat#4290, Hobbs Medical Inc, Stafford Springs, CT, USA) in the olfactory cleft under zero-degree endoscopic visualization. The brush was rotated briefly to collect surface mucosal cells. The brush was them submerged into collection solution [Hibernate A medium (Thermo Fisher, Waltham, MA)] with 5 μg/ml N-acetylcysteine (NAC, Sigma, St. Louis, MO, USA) on ice.

All biopsies were immediately digested as previously described^2^ for 15 minutes at 37°C with a 2.5 ml enzyme cocktail in Hibernate-A comprised of EDTA (1mM), Papain (2 mg/mL, Worthington Biochemical, Lakewood, NJ), DNAse I (StemCell Tech, Vancouver, BC, Canada) and NAC 5 μg/ml with frequent gentle trituration. After 15 minutes, an equal volume of 1x Accutase (StemCell Tech) was added, and samples were incubated for an additional 1 minute at 37°C. At the end of 1 minute, FBS (Thermo Fisher) was added. All samples were then filtered through a 70μm filter and centrifuged 5 minutes at 400 × g. Samples were washed in PBS with 2% FBS, spun and resuspended in HBSS or Hibernate-A containing 0.2% non-acetylated bovine serum albumin (Thermo), anti-clumping reagent 0.5 μl/ml (Gibco), and NAC 5 μg/ml (Sigma) to a final concentration of 1 million cells/mL.

### Single cell RNA-sequencing

Single-cell 3′ gene-expression matrices were generated with the 10x Genomics Chromium pipeline. Samples were processed for single cell analysis as described previously ^3^. Briefly, cells were quantified with a viability stain on an automated counter (Cellaca MX, Nexcelom) and loaded onto a Chromium iX controller (10X Genomics, Pleasanton, CA) for cell capture and bar coding targeting 20,000 cells, per the 3’ v4 gene expression protocol per manufacturer’s instructions. Reverse transcription, amplification, and library preparation were performed per protocol. Sequencing was performed at a depth of 300 million paired reads (NovaSeqX, Illumina). Each OE biopsy library was sequenced to a total depth of approximately 300 million paired-end reads (NovaSeq X, Illumina). Across olfactory epithelium samples, the mean sequencing depth was 57,062 reads per cell (range 30,849-131,047), with a median of 4,218 genes detected per cell (range 1,946-6,820). Single-cell 3′ sequencing data were demultiplexed and converted into FASTQ files using the 10X Genomics Cell Ranger v9.0.0 pipeline. Reads were aligned to the GRCh38-2024-A (Ensembl 98) human genome. Gene expression counts, barcodes, and UMIs were generated using cellranger count.

### Quality Control, normalization, and batch correction

Initial quality control, normalization, and batch correction were performed in Python (v3.9.13) using Scanpy (v1.9.1) following best practices for single cell analysis. Filtered feature matrices were processed as AnnData objects with sc.read_10x_mtx(), multiple scRNA-seq datasets were merged using sc.concat(join=”outer”) before analysis. Cells were excluded if they contained < 500 or > 8,000 detected genes, < 1,000 total UMIs or > 30 % mitochondrial transcripts. Normalization was performed with sc.pp.normalize_total() for a target sum of 10,000. Raw counts were copied into a dedicated *counts* layer. Library-size normalization to 10,000 transcripts per cell followed by natural-log transformation was stored in a *norm* layer for visualizations, preserving raw counts for statistical models.

Highly variable genes (HVGs) were defined with the Scanpy seurat_v3 flavor and refined with scvi.data.poisson_gene_selection. The subsetted AnnData object was registered with SCVI.setup_anndata using per-subject batch labels and the percentage of mitochondrial UMIs as a continuous covariate, and an SCVI model (negative-binomial likelihood, default 10-dimensional latent space) was trained on a single NVIDIA GPU for up to 500 epochs with early stopping (patience = 20). The learned latent representation was used to construct a k-nearest-neighbor graph, generate UMAP embeddings (min_dist = 0.5), and derive Leiden clusters. Cells for the initial Figure 1 cell atlas were labeled based on canonical gene markers (Extended Data Figure 1).

### Integration of CSF data

CSF data of 18 subjects (n = 9 control, n = 9 clinical AD) were obtained from the Gene Expression Omnibus (GEO) under accession number GSE134578 ^4^. Similar to concatenation above, filtered feature matrices were processed as AnnData objects with sc.read_10x_mtx(), multiple scRNA-seq datasets were merged using sc.concat(join=”outer”). Identical quality control, normalization, and batch correction was performed as above, except 5,000 HVGs were used, and “tissue source” (i.e., either OE or CSF) was used as an additional categorical variable in the ScVI model to correct for tissue source effects. Leiden clusters enriched for PTPRC, CD3D and CD3E were used to extract T-cell populations from both datasets; these cells were re-normalized, and the ScVI workflow was rerun on this T-cell subset for downstream analysis.

### T cell classification and analyses

T cell GEPs were obtained by querying the anndata against StarCAT (reference = TCAT.V1^5^), which contains GEPs from derived from cNMF of 1.7 million T cells from 700 individuals across 38 tissues. This database reproducibly classifies T cells using a logistic regression based on their dominant T cell identity GEP. T cell subset counts were aggregated per donor, converted to within-patient fractions for each source (CSF, OE) and AD stage (Control, Pre-Clinical, Clinical), averaged across patients in each group, and plotted as stacked bar charts. “Total memory” is the summed fraction of CD8-CM, CD8-EM, and CD8-TEMRA cells, whereas “CD4 vs CD8” is the ratio of pooled CD4 subsets (CM, EM, naïve, Treg) to pooled CD8 subsets (CM, EM, naïve, TEMRA).

Derived from the StarCAT reference, the ASA score integrates four gene-expression programs that were validated by demonstrating concordance between ASA-positive cells and activation-marker–defined populations in vivo, peptide-stimulated cells ex vivo, and clonally expanded T cells in silico, providing convergent evidence that ASA reliably marks TCR-driven, antigen-specific activation in humans^5^. For each subject, source, AD stage, and CD8 subset, we computed the percentage of ASA⁺ cells from replicate bulks of 10 cells (30 cells per patient). Group means and standard errors (mean ± SEM) were calculated across donors and displayed as bar charts. Donor-clustered binomial generalized linear models (GLMs) were used to test for stage-associated differences in ASA⁺ fractions within each source while controlling for individual subjects (OE: Control vs. Pre-Clinical, Control vs. Clinical; CSF: Control vs. Clinical).

### Myeloid cell classification and analyses

Myeloid cells were isolated by selecting the “Macrophage/DC” entry in the curated Figure 1 cell atlas, yielding 3,788 cells. Renormalization and ScVI pipeline was rerun on this subset similar to above with 10,000 HVGs. Cell-state identities were refined with StarCAT (reference = MYELOID.GLIOMA.V1 ^6^), which returned per-cell gene expression program (GEP)-usage scores for microglia, macrophage, monocyte, dendritic-cell and neutrophil identity. Rule-based thresholds (≥10 % program usage) assigned each cell to Microglia-like (≥ 10% use of macrophage and microglia GEP), Macrophage, Monocyte, cDC, or Neutrophil; the latter were excluded from downstream analyses. For each sample, we tallied the number of cells assigned to each myeloid category and expressed these counts as within-patient fractions. Fractions were then averaged across participants within each AD stage (Control, Pre-Clinical, Clinical) to obtain group-level means, which were visualized as a stacked bar plot (Extended Data Figure 3). Derived from the StarCAT reference, the four myeloid functional GEPs—Complement-Immunosuppressive, Scavenger-Immunosuppressive, Systemic-Inflammatory and Inflammatory-microglia—were shown by the authors to be bona-fide, plastic activation states by inducing and reversing them in patient-derived organoids, confirming the shifts via flow-cytometry, immunohistochemistry, and single-cell multi-omics^6^.

To model functional GEP usage, the four myeloid functional GEPs—Complement-Immunosuppressive, Scavenger-Immunosuppressive, Systemic-Inflammatory, and Inflammatory-microglia—were row-normalized so their weights summed to 1 per cell; a cell was deemed to have a dominant expression of that GEP when a single module accounted for > 50 % of its total usage. For each participant, we calculated the fraction of positive cells for every program, then averaged these fractions across subjects within the ordered disease stages (Control, Pre-Clinical, Clinical). Stage-wise means ± s.e.m. were visualized with faceted bar plots in Seaborn (Figure 3). Overall stage effects were assessed with a Kruskal–Wallis test; when significant, pair-wise differences (Control vs Pre-Clinical, Control vs Clinical, Pre-Clinical vs Clinical) were evaluated by two-sided Mann–Whitney U tests. Resulting *P*-values were multiplicity-corrected by Bonferroni adjustment.

### Myeloid cell pseudobulking, pathway enrichment, and differential expression analyses

For each donor with ≥45 myeloid cells, we created three non-overlapping pseudobulks by randomly aggregating 15 raw-count profiles per replicate (18 subjects × 3 replicates = 54 bulks, 1 subject had fewer than 45 myeloid cells); gene-wise sums were exported alongside bulk-level metadata. Differential expression was performed in edgeR: counts were TMM-normalized, a group-specific design matrix (Control, Pre-Clinical, Clinical) was fitted with the quasi-likelihood GLM pipeline, and contrasts were tested for Pre-Clinical vs Control and Clinical vs Control, retaining genes that passed filterByExpr. For pathway analysis, log-CPM values were analyzed with limma-fry against Human Phenotype Ontology (≥15 detected genes per set); brain-, CSF-, MRI- or neuro-related pathways were ranked by FDR and visualized as bar charts. Concordant and stage-specific DE genes were displayed in a double-sided quadrant volcano plot (log₂FC_Pre-Clinical on X, log₂FC_Clinical on Y; FDR < 0.05 threshold), and expression of exemplar genes was presented as stage-stratified box-and-whisker plots with pair-wise Wilcoxon tests (BH-adjusted).

### Olfactory sensory neuron classification and differential expression analysis

OSNs were isolated by selecting the “OSN” entry in the curated Figure 1 cell atlas. Renormalization and ScVI pipeline was rerun on this subset similar to above with 2,000 HVGs. OSNs were labeled based on Leiden clustering of dominant, canonical OSN subset genes. For each donor, 27 neurons were sampled by creating three non-overlapping pseudobulks with 9 OSNs. Donors contributing fewer than 27 OSNs in total were excluded to maintain uniform replication. The resulting gene-by-bulk matrix was analyzed in edgeR: low-expressed genes were filtered, libraries were TMM-normalized, tagwise dispersions were estimated, and quasi-likelihood GLMs were fit to test Pre-Clinical vs Control and Clinical vs Control contrasts, yielding FDR-adjusted DE tables for each comparison. Concordant and stage-specific DE genes were displayed in a double-sided quadrant volcano plot (log₂FC_Pre-Clinical on X, log₂FC_Clinical on Y; FDR < 0.05 threshold), and expression of exemplar genes was presented as stage-stratified box-and-whisker plots with pair-wise Wilcoxon tests (BH-adjusted).

### Cell communication modeling

T cells, myeloid cells, and OSNs were subject to renormalization and above ScVI pipeline with 10,000 HVGs. Single-cell transcriptomes from 22 donors were exported from Scanpy as an AnnData object and imported into R (v4.4.1). Raw UMI counts were transposed to a genes × cells dgCMatrix, per-cell library sizes were computed, and data were log-normalized to 10,000 counts/cell. Cell metadata included (i) donor identifiers (samples), (ii) Alzheimer’s stage (Control, Pre-Clinical, Clinical), and (iii) harmonized cell-type labels. For each stage a separate CellChat (v1.6.1). object was created, using the curated human protein-based CellChatDB (non-protein categories excluded). Following the standard workflow—subsetData, identifyOverExpressedGenes, identifyOverExpressedInteractions, computeCommunProb (trimean weighting, population-size correction off), and filterCommunication (≥10 source + target cells per pair)—we inferred ligand–receptor (LR) networks, computed pathway probabilities, aggregated signaling, and calculated centrality scores^7^.

To compare global network architecture, we merged stage objects (mergeCellChat) and contrasted either all three conditions simultaneously (compareInteractions) or pairwise (Control vs Pre-Clinical, Control vs Clinical). Differential LR usage between Clinical or Pre-Clinical AD compared to controls were identified using identifyOverExpressedGenes (logFC ≥ 0.05, pct-expressed ≥ 10 %, FDR ≤ 0.05), mapped back onto the network (netMappingDEG), and filtered to retain LR pairs up-regulated in the disease state (subsetCommunication). Gene-level chord diagrams (netVisual_chord_gene) were then generated to visualize directional changes in signalling: (i) mOSN + iOSN to all other cell types and (ii) reciprocal incoming signals, for both Pre-Clinical and Clinical stages.

### Olfactory epithelium biomarker analysis

Cells belonging to five broad lineages (Macrophage/DC, T cell, MV, neuron, sustentacular) were retained; scVI was used for reclustering and batch correction as above with 10,000 HVGs. Broad lineage labels were collapsed to Immune (macrophage, DC, and T cell) and Neuronal/Support (OSNS, MV, sustentacular). For each lineage we generated three non-overlapping pseudobulks of 75 immune or 30 neuronal/support clusters per donor by creating 3 bulks of 25 immune or 3 bulks of 10 neuronal/support cells. Pseudobulk counts were analyzed in edgeR v3.44 with quasi-likelihood GLMs. Libraries were TMM-normalized, low-expressed genes filtered by filterByExpr, and dispersions estimated with estimateDisp. Differential expression was tested for Pre-Clinical vs Control and Clinical vs Control contrasts; FDR-adjusted tables were exported for both lineages. Pseudobulk edgeR results for Immune and Neuronal/Support lineages (Pre-Clinical vs Control and Clinical vs Control contrasts) were imported into R (edgeR v3.44, ComplexHeatmap v2.18). For each lineage, genes with FDR < 0.05 and positive log₂-fold-change in either disease comparison were collected; the maximum logFC observed across all four tests was recorded, and non-informative loci (mitochondrial, histone, ENSG-prefixed or dot-containing symbols) were discarded. Lineage-specific pseudobulk count matrices and metadata were converted to log₂-CPM with a prior count of 0.5. Replicate bulks were averaged to a single expression profile for each donor × lineage combination, after which per-gene Z-scores were calculated. Genes showing positive Z-scores in every diseased sample of their lineage were retained, and the top 20 genes per lineage were chosen according to the previously calculated best logFC (40 genes total). Euclidean distances were hierarchically clustered (Ward’s method) and rows were split into two data-driven clusters. The final heat-map was rendered with ComplexHeatmap, coloring Z-scores from –2 (blue) to +2 (red) and annotating columns with Alzheimer’s stage, lineage, age, sex, APOE genotype, Aβ42/40 ratio and UPSIT score (bar-plots for continuous variables).

Module-level diagnostic performance was assessed at the patient level. For each donor we averaged per-gene Z-scores across the 20-gene Immune module, the 20-gene Neuronal-support module, and their sum (combined score). Patients were dichotomized (Control = 0; Pre-Clinical + Clinical AD = 1), and empirical ROC curves together with point AUCs were generated with pROC (roc). Ninety-five-percent confidence intervals for each AUC were estimated by 2,000-fold stratified bootstrap resampling. Curves were plotted with a diagonal reference line, and subplot subtitles report the AUC and its bootstrap CI for the Immune, Neuronal-Support, and Combined scores. Pearson correlation coefficients (R) and corresponding two-tailed p values between the Combined ΔZ score and each clinical or biomarker variable were calculated with the stat_cor function in ggpubr (method = “pearson”), and the resulting R and p values are reported in the figure panels.

### Flow Cytometry

Olfactory Epithelium (OE) biopsies were thawed in a 37ªC water bath from liquid nitrogen frozen in 90% FBS and 10% DMSO. For each centrifugation step, 400 x *g* for 5 minutes at 4ªC was used. For each AD status, 3 biological replicates each with 3 technical replicates were stained and analyzed. Cells were washed twice with dPBS, then stained with viability aqua live-dead stain in dPBS (1:200, Thermo Fisher) for 30 minutes. Cells were resuspended in FACS buffer (2% FBS, 2mM EDTA in PBS) with CD16 and CD32 with 2% normal human serum for 20 minutes at 4ªC for non-specific antibody blocking. Cells were then washed with FACS buffer, and resuspended in antibody cocktail (FITC-anti-human HLA-DR, Clone: LN3, Cat: 327005, RRID: AB_893577 BV745/750-anti-human CD3, Clone: SK7, Cat: 344845, RRID: AB_2734352 APC/Cy7 anti-human CD8a, Clone: HIT8a, Cat: 300925, RRID: AB_10612924 PE anti-human CD38, Clone HIT2, Cat: 303505, RRID: AB_314357, all at 1:100 and from Biolegend) for 30 minutes. Cells were washed and fixed in 0.8% PFA. Compensation beads and OE cells with single color-antibody controls were used for spectral compensation. Fluorescent-minus one (FMO) samples were used to identify CD38 and HLA-DR positivity from background. All samples were immediately analyzed using an LSRFortessa Cell Analyzer (BD) and analyses were done in FlowJo v10.10.

### Statistical Analyses

Twenty two donors were analyzed, cohort size was chosen to match typical discovery single-cell RNA-seq studies and to obtain similar numbers per disease stage. No formal power calculation was performed; pilot down-sampling confirmed that the principal cell-state and gene-expression differences were stable at this sample size. There were no data exclusions. Results were declared significant at a false-discovery rate (FDR) < 0.05 or a family-wise–error–corrected P < 0.05. Exact biological replicate numbers are provided in figure legends. Group differences in cell-type proportions were evaluated with Kruskal–Wallis tests followed, when the omnibus test was significant, by pairwise two-sided Mann–Whitney U tests with Bonferroni correction; two-group comparisons in CSF (Control vs. Clinical) used Wilcoxon rank-sum tests. For analyses of ASA-positive fractions across CD8 memory subsets, donor-clustered binomial generalized linear models (GLMs) were used. Each donor-level 30 cell replicate was modeled as a binomial outcome representing the number of ASA-positive cells out of the total cells per replicate. Models included AD stage as a categorical fixed effect and accounted for within-donor correlation using cluster-robust standard errors. Pre-specified pairwise contrasts were computed within each source (OE: Control vs. Pre-Clinical and Control vs. Clinical; CSF: Control vs. Clinical). Pseudobulk differential expression employed edgeR quasi-likelihood generalized linear models, reporting log₂-fold-changes, quasi-likelihood F statistics with corresponding degrees of freedom, raw P values and Benjamini–Hochberg FDR in supplemental tables. For pathway enrichment, limma-fry was applied to log-CPM values and set-level P values were FDR-adjusted. Diagnostic performance of the 20-gene Immune module, 20-gene Neuronal-Support module and their Combined score was assessed with receiver-operating-characteristic (ROC) analysis: empirical ROC curves and point areas under the curve (AUCs), 95 % confidence intervals were estimated from 2,000 stratified bootstrap replicates. Figures display mean ± s.e.m. for bar charts and median with 25–75 % inter-quartile range for box-and-whisker plots. Pearson correlation coefficients (R) and corresponding two-tailed p values between the Combined ΔZ score and each clinical or biomarker variable were calculated and reported in the figure panels.

## Code Availability

All analyses for scRNA-seq were performed with publicly available software packages. Code for reproducing results will be available at Goldstein github (https://github.com/Goldstein-Lab) and is available upon request.

## Extended Data

**Extended Data Figure 1.**
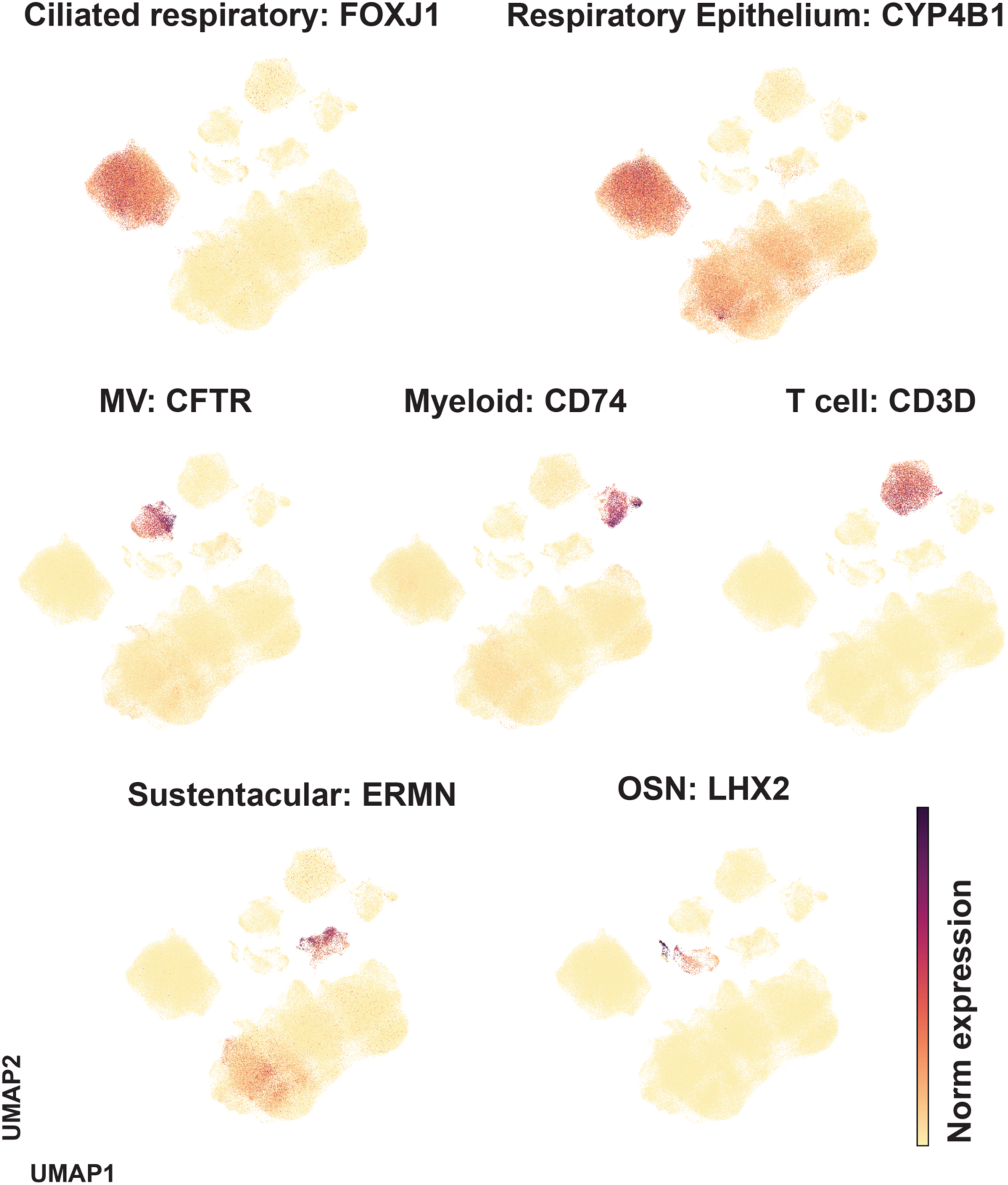
Example genes from OE cell atlas. UMAP shows one exemplary gene from each of the 7 major cell subsets from Figure 1 including ciliated respiratory, respiratory epithelium, microvillar cells (MV), myeloid cells, T cells, sustentacular cells, and olfactory sensory neurons (OSN). Dark red color represents a higher normalized gene expression.

**Extended Data Figure 2.**
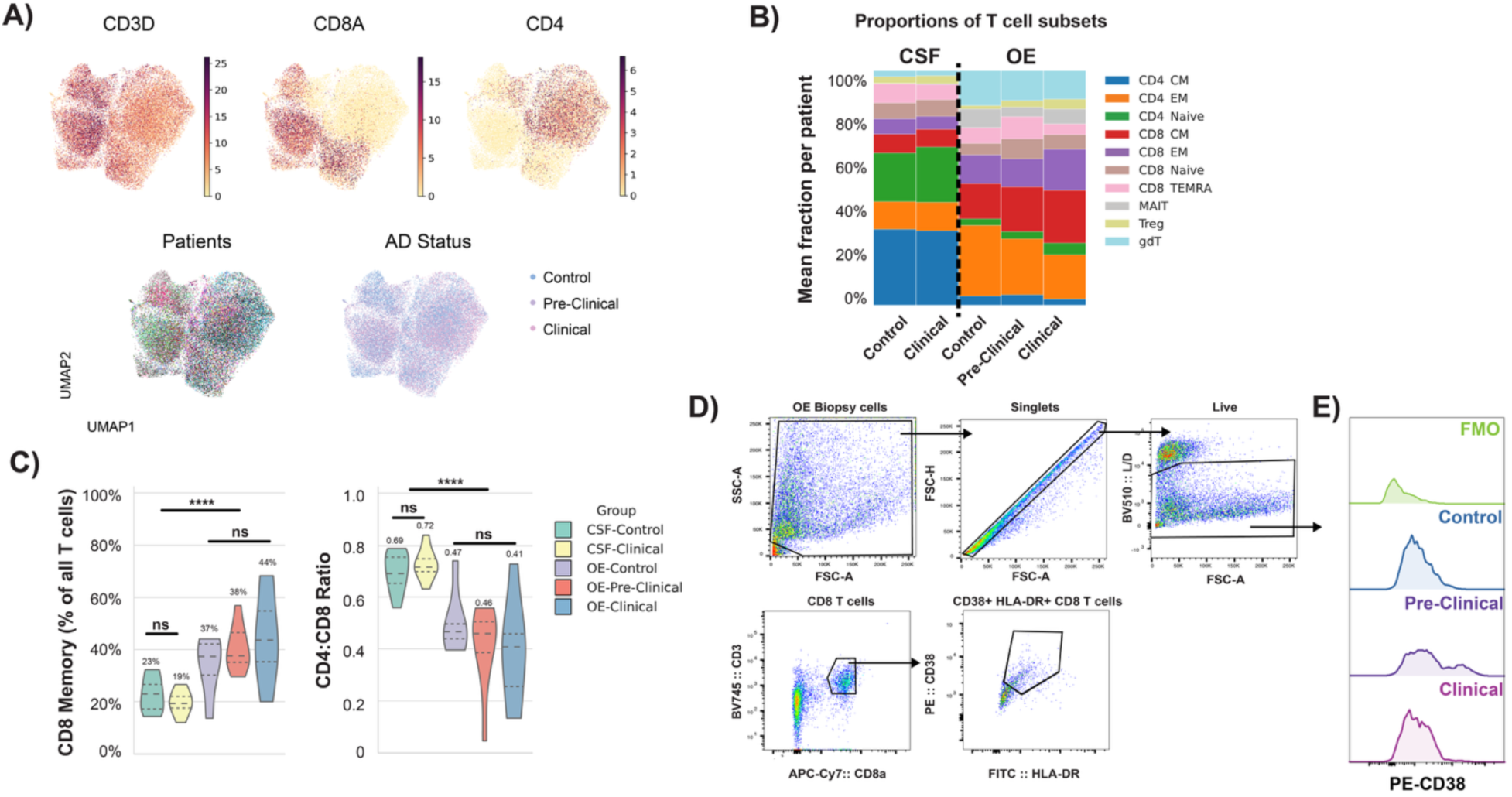
T cell gene expression and flow cytometry analysis. **A)** UMAPs show canonical T cell markers, CD3D, CD8A, and CD4 across OE and CSF T cells (top), or by individual subjects and AD status (bottom). **B)** Graph of mean proportion of each T cell subset per subject by source and AD stage (Treg = regulatory T cell; gdT = gamma delta T cell). **C)** Violin plots show subject-level T-cell metrics CSF and OE across Alzheimer’s disease stages. The left panel (“Total memory”) gives the summed fraction of memory CD8⁺ subsets (CM + EM + TEMRA) within each sample, while the right panel shows the ratio of CD4 (CM + EM + naïve + Treg) to CD8 (CM + EM + naïve + TEMRA) cells. Violins outline median and quartile distribution, overlaid significance bars indicate Mann-Whitney U (CSF Control vs Clinical), Kruskal–Wallis (OE Control, Pre-Clinical, Clinical), and Mann-Whitney U (CSF vs OE) tests, with *** p ≤ 0.001. **D)** Shows representative gating strategy of CD38+ HLA-DR+ CD8+ T cells from the OE brush biopsies. **E)** Representative histograms show modal expression of CD38 in fluorescent-minus-one (FMO) control, as well as in Control, Pre-Clinical, and Clinical AD subjects.

**Extended Data Figure 3.**
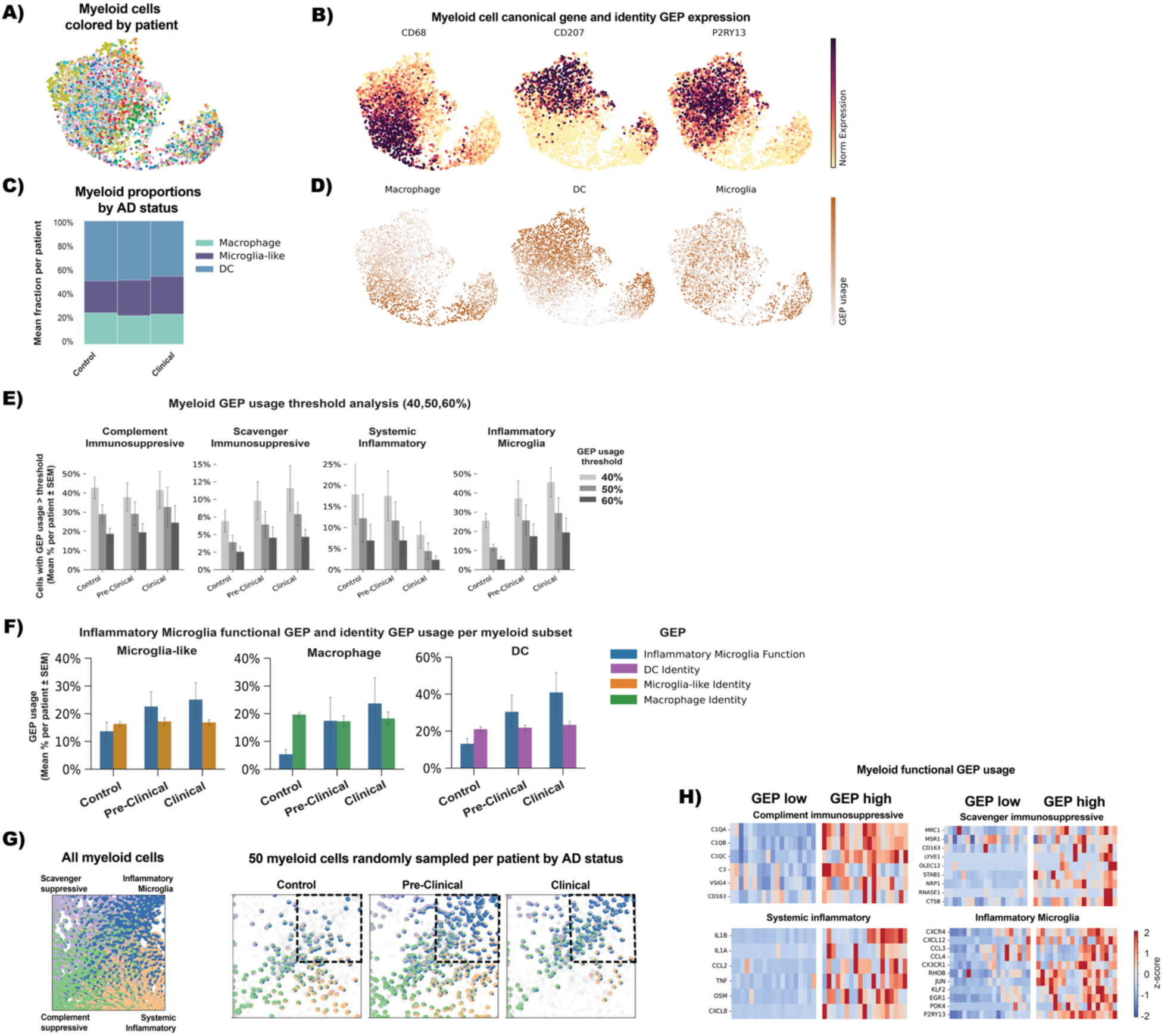
Myeloid characterization and function. **A)** UMAP shows all myeloid cells colored by individual subject. **B)** same UMAP overlayed with canonical myeloid genes including macrophage (CD68; left), dendritic cell (CD207; middle) and microglia-like (P2RY13; right). **C)** shows the mean fraction of myeloid cells among all myeloid cells per subject by AD status. **D)** UMAPs overlaid with myeloid GEP scores used for classification. **E)** Myeloid GEP usage threshold analysis was performed at 40%, 50%, and 60%. The proportion of myeloid cells per patient exceeding each GEP usage threshold is shown for four modules: Complement Immunosuppressive, Scavenger Immunosuppressive, Systemic Inflammatory, and Inflammatory Microglia. Bars indicate mean ± SEM across patients. Data shows that GEP thresholds do not change data interpretation. **F)** Mean percentage of cells per patient using each identity GEP within microglia-like, macrophage, or dendritic cell (DC) subsets and inflammatory microglia function module. Data shows that cell identity is constant throughout AD groups, but inflammatory microglia function changes across groups, consistent across cell types. **G)** Quadrant plot of individual cells with each of the four functional GEPs (scavenger suppressive, inflammatory microglia, complement suppressive, systemic inflammatory) represented in each corner, where coordinates show relative abundance of each GEP. Right plots are the same quadrant plot as E) but with 50 myeloid cells randomly sampled per subject by AD status, shows overrepresentation of inflammatory microglia GEP in the preclinical and clinical AD stages. **H)** Heatmaps show GEP low (<10% usage) and GEP high (>10% usage) z-scored expression of top genes for each program (rows) by subject (columns).

**Extended Data Figure 4.**
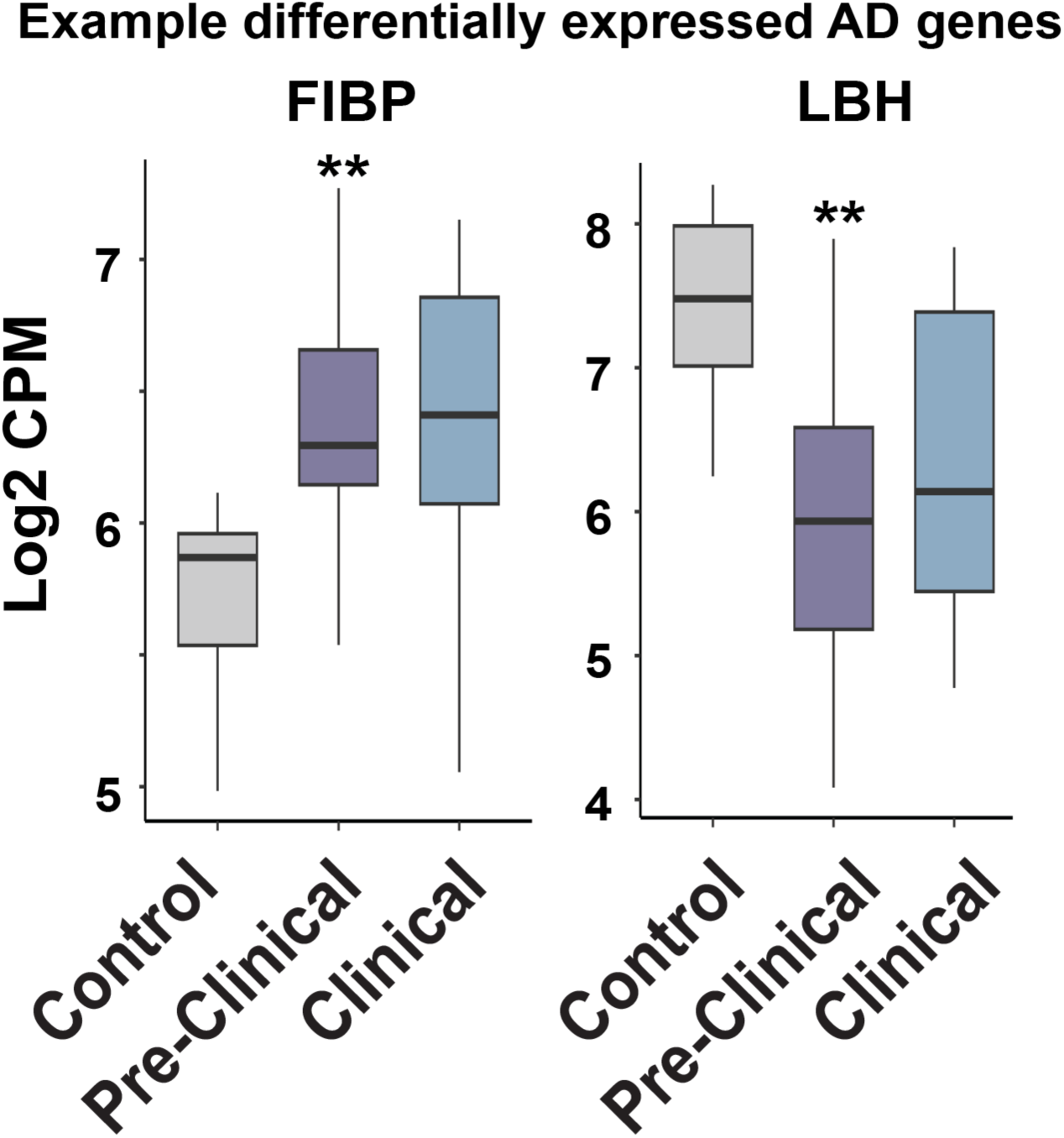
Example AD associated differentially expressed OSN genes. Boxplots show donor-level pseudobulk log₂ counts per million (CPM) for FIBP and LBH across Control, Pre-Clinical, and Clinical AD groups. Boxes represent the median and interquartile range. Significance was assessed by pairwise two-sided Wilcoxon rank-sum tests with Benjamini–Hochberg correction; p < 0.01.

## Supplementary Data Legends

**Supplemental Video 1. Endoscopic olfactory cleft brush biopsy technique.** Video shows a 0-degree nasal endoscopy view of a cytology brush biopsy of the right olfactory cleft in an awake subject in the outpatient clinic. The nasal cavity was previously decongested and anesthetized using topical oxymetazoline and tetracaine. The biopsy portion of the cytology brush is positioned between the middle turbinate and the nasal septum and carefully advanced to contact the superior turbinate laterally and the nasal septum medially and then rotated to sample the surface epithelium of the olfactory cleft. Effort is made to position the brush superiorly and to minimize brush contact with non-olfactory regions.

**Table S1 Myeloid_DE_preclinical_vs_control.csv** Differential-expression output from the edgeR quasi-likelihood test comparing pre-clinical AD pseudobulks from myeloid cells with controls; each row is a retained gene and columns report log₂ fold-change (Pre-Clinical – Control), average log-CPM, quasi-likelihood F-statistic, raw P-value, and Benjamini–Hochberg FDR.

**Table S2 Myeloid_DE_clinical_vs_control.csv.** Identical format to S1, but the contrast is clinical AD versus control.

**Table S3 OSN_DE_preclinical_vs_control.csv** Differential-expression output from the edgeR quasi-likelihood test comparing pre-clinical AD pseudobulks from OSNs with controls; each row is a retained gene and columns report log₂ fold-change (Pre-Clinical – Control), average log-CPM, quasi-likelihood F-statistic, raw P-value, and Benjamini–Hochberg FDR.

**Table S4 OSN_DE_clinical_vs_control.csv.** Identical format to S4, but the contrast is clinical AD versus control.

**Table S5 Combined_module_deltaZ_by_patient.csv.** Table lists each subject’s AD status alongside the Δ-Z score used in the final “Combined Module” box-and-whisker plot in Figure 5C.

**Table S6.** Immune and Neuronal-Support gene modules enriched in Alzheimer’s disease. The table lists the top 20 candidate genes per module that were significantly up-regulated (log₂ FC > 0, FDR < 0.05) in either pre-clinical or clinical AD pseudobulks. Genes were selected after filtering out mitochondrial, histone, and unannotated loci and ranked by their maximum positive log₂ fold-change across comparisons. The “module” column denotes assignment to the Immune or Neuronal/Support clusters.

| Gene | Module |
| --- | --- |
| ARHGAP45 | Immune |
| CCDC88B | Immune |
| CD6 | Immune |
| CYTOR | Immune |
| FLNA | Immune |
| GZMK | Immune |
| HLA-DQA2 | Immune |
| HSPA6 | Immune |
| IL12RB1 | Immune |
| ITGB2 | Immune |
| LCK | Immune |
| LINC01943 | Immune |
| MATK | Immune |
| MFNG | Immune |
| MYO1G | Immune |
| S100B | Immune |
| TBC1D10C | Immune |
| TMC8 | Immune |
| TNNI2 | Immune |
| ZAP70 | Immune |
| C19orf33 | Neuronal_Support |
| C6orf52 | Neuronal_Support |
| CA11 | Neuronal_Support |
| CDC37L1-DT | Neuronal_Support |
| CYP26A1 | Neuronal_Support |
| DMWD | Neuronal_Support |
| FAM83H | Neuronal_Support |
| FUT2 | Neuronal_Support |
| LGALS3BP | Neuronal_Support |
| LRRC24 | Neuronal_Support |
| MAPK8IP1 | Neuronal_Support |
| MISP | Neuronal_Support |
| NR2F6 | Neuronal_Support |
| PLEKHH3 | Neuronal_Support |
| S100A14 | Neuronal_Support |
| SCRIB | Neuronal_Support |
| SYCE1L | Neuronal_Support |
| THNSL2 | Neuronal_Support |
| TUBB2B | Neuronal_Support |
| VWA1 | Neuronal_Support |

